# Foreign Body Responses Adapt to Systemic Immunosuppression to Maintain Local Immune Homeostasis Through Decorin-Associated Signaling

**DOI:** 10.64898/2026.10.27.762341

**Authors:** Yuwei Zhang, Jing Wang

## Abstract

Foreign body responses (FBRs) involve complex interactions between pro- and anti-inflammatory signals that regulate the integration and function of implanted biomaterials. However, most mechanistic studies of FBRs have been conducted in healthy hosts, and little is known about how these immune responses are regulated under pathological conditions. Here, we investigated immune responses to porous polycaprolactone (PCL) scaffold implants in mice with endotoxin tolerance or aggressive breast tumors, which induce systemic immunosuppression through distinct mechanisms. Despite suppressed inflammatory activity in major organs, implanted scaffolds maintained locally immune-active environments characterized by increased pro-inflammatory immune-cell populations and elevated inflammatory mediators, including CXCL1. In migration assays, CXCL1 preferentially recruited neutrophils and enriched immune-cell populations with pro-inflammatory phenotypes among splenocytes derived from both endotoxin-tolerant and tumor-bearing mice. The extracellular matrix proteoglycan decorin, a damage-associated molecular pattern with TLR2/4-activating activity in myeloid cells, was enriched within scaffolds but reduced in major organs under both immunosuppressive conditions. Recombinant decorin promoted pro-inflammatory phenotypes in suppressive myeloid-cell populations and enhanced CXCL1 production through TLR2/4-associated signaling, suggesting a mechanistic link between decorin-mediated myeloid activation, CXCL1-dependent immune-cell recruitment, and the maintenance of local immune activity at biomaterial interfaces. Together, these findings reveal that FBRs are not merely passive inflammatory reactions to biomaterial implants but can dynamically adapt to systemic immune dysfunction to sustain local immune homeostasis. This intrinsic immunoregulatory capacity may represent an additional factor influencing the performance of immunomodulatory implants and should be considered in biomaterial design to optimize therapeutic outcomes across different applications and host immune conditions.

**Graphic abstract:** 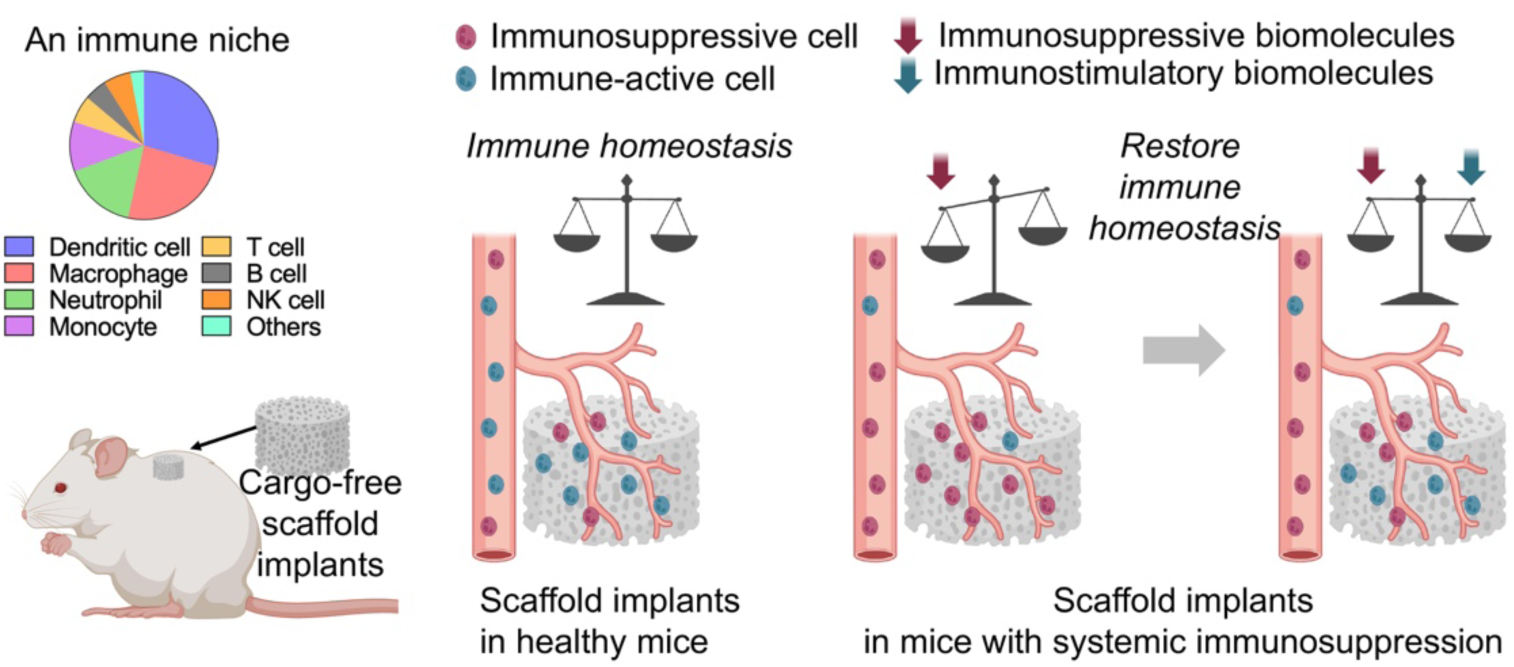

## Introduction

Biomaterial scaffold implants are widely used as platforms for tissue engineering and for the localized delivery of therapeutic drugs, biologics, and cells. Following implantation, scaffolds elicit a host immune response known as foreign body response (FBR) [1; 2; 3; 4]. The FBR begins with an acute phase characterized by protein adsorption, provisional matrix formation, and recruitment of neutrophils and macrophages. This response later progresses into a chronic phase marked by macrophage fusion into foreign body giant cells, fibroblast recruitment, angiogenesis, extracellular matrix deposition and remodeling, and, in some cases, fibrous capsule formation [1; 2; 4; 5]. These processes are strongly influenced by scaffold properties, including biomaterial composition, surface chemistry, porosity and pore size, stiffness, and degradation products [3; 6; 7; 8; 9].

The FBR has traditionally been viewed as a pathological inflammatory reaction that compromises implant function. However, it can also be considered a dynamic immune-remodeling process that attempts to restore local tissue homeostasis [3; 10; 11]. Inflammatory signals are required to clear debris, prevent infection, and initiate repair, whereas anti-inflammatory and pro-resolving signals limit excessive tissue damage and prevent uncontrolled fibrotic encapsulation. Although many studies have examined how different biomaterials and scaffold architectures induce varying degrees of FBR, most have been performed in healthy hosts. Much less is known about how an established FBR adapts when disease disrupts systemic immune homeostasis and disease-associated inflammatory or immunosuppressive cells infiltrate the scaffold environment.

In our previous work, we compared the immune environments within porous polycaprolactone (PCL) scaffolds implanted in healthy mice and in mice bearing aggressive breast tumors, which develop progressive systemic immunosuppression [12]. PCL is a biocompatible, slowly degradable polymer widely used in tissue engineering and regenerative medicine. Porous PCL scaffolds generally elicit a localized and relatively mild FBR, characterized by immune- and stromal-cell infiltration, vascularization, and tissue integration [13; 14; 15]. In tumor-bearing mice, cancer-induced suppressive myeloid cells accumulated in both major organs and scaffold implants. These cells secreted cytokines and chemokines that promoted the recruitment of circulating tumor cells to both sites. However, major organs and scaffolds responded differently to this infiltration. Major organs progressively developed significant immunosuppressive environments that supported metastatic outgrowth. In contrast, scaffolds maintained an immune-active environment, which counteracted suppressive signals derived from suppressive myeloid cells and limited secondary tumor growth [12]. These findings suggested a previously underrecognized feature of the FBR: its potential to preserve localized immune homeostasis despite systemic immune dysregulation.

In the present study, we sought to further establish this homeostatic feature of the FBR across distinct pathological conditions and elucidate the underlying mechanisms by which implant-associated immune environments resist systemic immunosuppression. To this end, we investigated porous PCL scaffolds in mice bearing aggressive breast tumors and mice with lipopolysaccharide (LPS)-induced endotoxin tolerance, two models that develop systemic immune suppression through distinct mechanisms. Cancer-induced immunosuppression develops progressively in response to tumor-derived cytokines, antigens, metabolites, and other soluble factors. It is characterized by the expansion of suppressive myeloid populations, T-cell dysfunction, and elevated production of immunosuppressive mediators, including TGF-β and IL-10 [16]. In contrast, endotoxin tolerance is characterized by innate immune reprogramming following repeated exposure to LPS, during which macrophages and other innate immune cells become hyporesponsive to subsequent Toll-like receptor 4 (TLR4) stimulation and exhibit reduced production of pro-inflammatory cytokines, including TNF-α, IL-1β, and IL-12. This state can also be accompanied by impaired dendritic-cell maturation and enhanced regulatory immune activity, including enrichment of regulatory T cells (Tregs) in major organs [17; 18]. By examining these two mechanistically distinct models, we aimed to determine whether the preservation of implant-associated immune activity represents a generalizable feature of the FBR rather than a response specific to cancer-induced immunosuppression.

To investigate this possibility, we first subcutaneously implanted porous PCL scaffolds in immunocompetent BALB/c mice to establish an implant-associated FBR under healthy conditions. We subsequently induced endotoxin tolerance through repeated LPS administration or established aggressive breast tumors through inoculation of 4T1 murine breast cancer cells at the mammary fat pad. Following the development of systemic immunosuppression, we characterized and compared immune-cell populations and cytokine profiles within implanted scaffolds and major organs from healthy control, LPS-tolerant, and tumor-bearing mice. These analyses identified decorin, a small leucine-rich proteoglycan and extracellular matrix component with TLR2/4-agonistic activity, as a potential mediator of implant-associated immune homeostasis. Notably, decorin was enriched within scaffolds but decreased in major organs following the induction of systemic immunosuppression. Previous studies have primarily implicated decorin in extracellular matrix remodeling and fibrotic responses associated with biomaterial implantation [19; 20]. Additionally, decorin has been identified as an endogenous damage-associated molecular pattern (DAMP) that activates TLR2/4 signaling and exacerbates inflammatory disease progression [21; 22; 23; 24; 25; 26]. Building on these observations, we investigated how decorin interacts with disease-associated suppressive myeloid cells and contributes to the maintenance of local immune activity within the scaffold environment. Our findings reveal a previously underappreciated role of the FBR in sustaining localized immune homeostasis under systemic immunosuppressive conditions and identify decorin-associated innate immune signaling as a potential mechanism underlying this response. These findings advance our understanding of host–biomaterial interactions and provide insights for designing immunomodulatory implants, as the immune-homeostatic properties of the FBR may either enhance or limit the intended immunoregulatory effects of these implants.

## Materials and Methods

### Materials

Polycaprolactone (PCL, Mw 80,000) was from Polysciences. Poly(vinyl alcohol) (PVA) was from Thermo Fisher Scientific. Lipopolysaccharide (LPS) was from MilliporeSigma. Recombinant mouse CXCL1 was from BioLegend. Recombinant human decorin proteins and Alexa Fluor 647-conjugated mouse CXCL1 antibodies were from R&D systems. All other antibodies for flow cytometric analysis were BioLegend. OxPAPC, anti-mouse TLR2 antibodies (clone T2.5), and anti-mouse TLR4 antibodies (clone MTS510) were from InvivoGen. Unless otherwise specified, all other reagents were from Fisher Scientific.

### Cell culture

The 4T1/Luc/tdTomato murine breast cancer cells were kindly provided by Dr. Lonnie Shea at the University of Michigan. These 4T1 cells were cultured in high-glucose Dulbecco’s Modified Eagle’s Medium (DMEM) supplemented with 10% fetal bovine serum (FBS), 1% penicillin–streptomycin, 1% nonessential amino acids, 2 μg/mL puromycin and 300 μg/mL G418. Cells were maintained at 37°C in a humidified incubator with 5% CO₂.

### PCL scaffold fabrication

Porous PCL scaffolds were fabricated as described previously [12]. Briefly, a 6% PCL solution was homogenized with a 10% PVA solution at a volume ratio of 3: 20 and subsequently added to ultrapure water. The mixture was stirred overnight to facilitate microsphere formation, and the resulting microspheres were washed six times with ultrapure water. Following lyophilization for 48 h, the PCL–PVA microspheres were mixed with sodium chloride particles (250–425 μm) at a mass ratio of 1: 30. A total of 100 mg of the mixture was transferred into a cylindrical mold and compressed to form scaffolds measuring 8 mm in diameter and 3 mm in height. Both surfaces of the compressed scaffold–salt mixture were heated at 85 °C for 10 min each to promote microsphere fusion. The scaffolds were then immersed in deionized water at room temperature for 48 h to remove the salt particles and generate a porous structure. Finally, the scaffolds were sterilized with 70% ethanol, rinsed with ultrapure water, blotted dry using sterile absorbent paper, and stored at −80 °C until use.

### Scaffold implantation, LPS tolerance induction, and tumor inoculation in animals

Porous PCL scaffolds were implanted subcutaneously in the upper dorsal region of 7–8-week-old female BALB/c mice (the Jackson Laboratory), as previously described [12]. Two weeks after scaffold implantation, mice were subjected to repeated low-dose LPS administration to induce endotoxin tolerance or inoculated with 4T1 breast cancer cells to establish an aggressive tumor model. To induce endotoxin tolerance, LPS prepared in sterile PBS was administered intraperitoneally once daily using an escalating-dose regimen: 0.25 mg/kg for 2 days, 0.5 mg/kg for 2 days, 1 mg/kg for 2 days, 2.5 mg/kg for 3 days, and 5 mg/kg for 9 days [27]. Control mice received an equivalent volume of sterile PBS on the same schedule. Mice were euthanized 1 day after the final administration, and the scaffold, lung, spleen, and liver were collected for downstream analyses. To establish an aggressive breast cancer model, 1×10^6^ of 4T1 cells suspended in 100 µL of sterile PBS were inoculated into the mammary fat pad. Mice were euthanized when primary tumors reached ∼1,000 mm³, and the implanted scaffolds and lungs were harvested for subsequent analyses [12]. Each group included three mice (n = 3). All animal procedures were approved by the Iowa State University Institutional Animal Care and Use Committee and conducted in accordance with approved institutional guidelines and protocols.

### Tissue processing and immune cell profiling with flow cytometry

Scaffolds and major organs were minced into 2–3-mm pieces and enzymatically digested with 0.1 mg/mL Liberase (Roche) for 30 min at 37°C. The digested tissues were then passed through 70-μm cell strainers to generate single-cell suspensions. Red blood cells were removed using Red Blood Cell lysis buffer according to the manufacturer’s instructions (BioLegend). The resulting cells were resuspended in MACS buffer consisting of PBS supplemented with 0.5% bovine serum albumin and 2 mM EDTA. Cells were first blocked with anti-mouse CD16/32 antibody for 15 min and then stained with antibody cocktails against cell-surface markers for 30 min on ice. Subsequently, cells were fixed and permeabilized using Fixation/Permeabilization buffer (eBioscience™) and stained with antibody cocktails against intracellular markers overnight at 4°C. The following day, cells were washed with MACS buffer once and then analyzed using a BD FACSCanto flow cytometer. Flow-cytometry data were analyzed with FlowJo software (BD Biosciences) following the procedure described in our previous publication [28].

A total of five antibody panels were used to characterize immune cell populations. Panel 1 was designed to identify T helper cells (CD45⁺CD3⁺CD4⁺) and their major subsets, including Th1 cells (IFN-γ⁺), Th2 cells (IL-4⁺), and Th17 cells (IL-17A⁺). This panel included APC-Cy7-conjugated anti-CD45 (clone 30-F11), Pacific Blue-conjugated anti-CD3 (clone 17A2), BV605-conjugated anti-CD4 (clone GK1.5), PE-conjugated anti-IFN-γ (clone XMG1.2), APC-conjugated anti-IL-4 (clone 11B11), and PE-Cy7-conjugated anti-IL-17A (clone TC11-18H10.1) antibodies. Panel 2 was designed to identify regulatory T cells (Tregs), defined as CD45⁺CD3⁺CD4⁺CD25⁺Foxp3⁺ cells. This panel included APC-Cy7-conjugated anti-CD45, Pacific Blue-conjugated anti-CD3, BV605-conjugated anti-CD4, PE-conjugated anti-CD25 (clone PC61), and Alexa Fluor 488-conjugated anti-Foxp3 (clone MF-14) antibodies. Panel 3 was designed to identify Gr-1⁺ myeloid cells (CD45⁺CD11b⁺Gr-1⁺), with neutrophils (Ly6G⁺) and monocyte-like cells (Ly6G⁻) defined as subsets of this population. Each subset was further characterized according to IL-12/IL-23p40 and IL-10 expression. This panel included APC-Cy7-conjugated anti-CD45, BV711-conjugated anti-CD11b (clone M1/70), Pacific Blue-conjugated anti-Gr-1 (clone RB6-8C5), Alexa Fluor 488-conjugated anti-Ly6G (clone 1A8), APC-conjugated anti-IL-10 (clone JES5-16E3), and PE-conjugated anti-IL-12/IL-23 p40 (clone C15.6) antibodies. Panel 4 was designed to identify macrophages (CD45⁺CD11b⁺F4/80⁺) and to distinguish their subsets according to IL-12 and IL-10 expression. This panel included APC-Cy7-conjugated anti-CD45, BV711-conjugated anti-CD11b, Pacific Blue-conjugated anti-F4/80 (clone BM8), APC-conjugated anti-IL-10, and PE-conjugated anti-IL-12/IL-23 p40 antibodies. Panel 5 was designed to identify dendritic cells (CD45⁺CD11c⁺F4/80⁻) and their mature subset (CD86⁺MHC II⁺). This panel included APC-Cy7-conjugated anti-CD45, BV605-conjugated anti-CD11c (clone N418), Pacific Blue-conjugated anti-F4/80, PE-conjugated anti-CD86 (clone GL-1), and Alexa Fluor 488-conjugated anti-MHC II antibodies (clone M5/114.15.2).

### Cytokine array and ELISA

Conditioned media were prepared by incubating tissue-derived single-cell suspensions at a density of 5 × 10^6^ cells/mL in serum-free, phenol red-free RPMI 1640 medium for 24 h at 37°C. Following incubation, the conditioned media were collected and centrifuged at 500 × g for 5 min to remove cells and debris. Total protein concentrations were quantified using the BCA protein assay kit according to the manufacturer’s instructions (Thermo Fisher Scientific). Before analysis, all samples were diluted to a final total protein concentration of 500 μg/mL. Cytokine profiles in the conditioned media were analyzed using the Mouse Cytokine Array C3 membrane according to the manufacturer’s instructions (RayBiotech). Chemiluminescent signals were captured using an iBright 1500 Imaging System (Thermo Fisher Scientific). Spot intensities were quantified using iBright Analysis Software (Thermo Fisher Scientific) and normalized to the internal positive-control spots on each membrane. The final data were presented as normalized cytokine or protein levels. Additionally, decorin concentrations in the conditioned media were quantified using a mouse decorin ELISA kit according to the manufacturer’s instructions (RayBiotech).

### Transwell cell migration assay

The migratory responses of splenocytes were evaluated using a Boyden chamber Transwell assay. The spleens of LPS-tolerant or 4T1 tumor-bearing BALB/c mice were prepared to single-cell suspensions and resuspended in serum-free RPMI 1640 medium. A total of 5 × 10⁵ cells were added to the upper chambers of Transwell inserts with 8-μm pores. The lower chambers contained serum-free RPMI 1640 medium supplemented with 50 ng/mL recombinant CXCL1. Cells were incubated for 24 h at 37°C in a humidified incubator containing 5% CO₂. Splenocytes that migrated into the lower chambers and aliquots of the unsorted splenocytes were analyzed by flow cytometry.

### Suppressive Ly6G^+^ neutrophil isolation and their incubation with decorin

Lungs were collected from BALB/c mice bearing 4T1 tumors of ∼1,000 mm³ and processed into single-cell suspensions as described above. Ly6G⁺ neutrophils were isolated using magnetic-activated cell sorting (MACS) with a mouse neutrophil isolation kit according to the manufacturer’s instructions (Miltenyi Biotec). Isolated neutrophils were divided into four groups: (1) untreated control, (2) decorin treatment (100 ng/mL), (3) decorin treatment with OxPAPC, and (4) decorin treatment with anti-TLR2 and anti-TLR4 blocking antibodies. OxPAPC, an inhibitor of TLR2/4 signaling, and TLR2/4 blocking antibodies were used to assess the involvement of these receptors in decorin-mediated responses. For TLR2/4 inhibition studies, cells were preincubated with OxPAPC (30 µg/mL) or anti-TLR2 and anti-TLR4 blocking antibodies (5 µg/mL each) for 2 h, washed with PBS, and subsequently treated with decorin (100 ng/mL) for 24 h.

Cells and conditioned media were subsequently collected for flow cytometric and cytokine array analyses, respectively, following the procedures described above. Two antibody panels were used for flow cytometry. Panel 1 included PE-conjugated anti-phospho-NF-κB p65 (Ser529) and Alexa Fluor 488-conjugated anti-phospho-p38 MAPK (Thr180/Tyr182) antibodies. Panel 2 included Alexa Fluor 647-conjugated anti-mouse CXCL1 and PE-conjugated anti-IL-12/IL-23 p40 antibodies. Conditioned media were analyzed using the Mouse Cytokine Array C3.

### Statistical analysis

Data are presented as the mean ± standard error of the mean (SEM). Each experiment was independently repeated two to three times, with three replicates per sample. Statistical analyses were performed using GraphPad Prism. Comparisons between two independent groups were performed using an unpaired, two-tailed Student’s *t*-test. A p value of < 0.05 was considered statistically significant.

## Results

### PCL scaffold implants maintain local immune activity in LPS-tolerant and 4T1 tumor-bearing mice despite systemic immunosuppression

Our previous study demonstrated that 4T1 tumor-bearing mice developed systemic immunosuppression when primary tumors reached ∼1,000 mm³. Despite this systemic immune dysfunction, implanted scaffolds maintained a more immune-active environment than the immunosuppressive lung and other major organs, as evidenced by higher frequencies of pro-inflammatory immune-cell subsets and elevated levels of pro-inflammatory cytokines [12]. To determine whether this phenomenon extends beyond cancer-induced immunosuppression, we further investigated scaffold-associated immune responses in mice with LPS-induced endotoxin tolerance, a mechanistically distinct form of systemic immune dysfunction.

Porous PCL scaffolds measuring 8 mm in diameter and 3 mm in height were subcutaneously implanted in the dorsal flank of immunocompetent BALB/c mice. Two weeks after implantation, when wound healing, scaffold vascularization, and cellular infiltration were largely established, mice received repeated intraperitoneal injections of LPS at escalating doses over 18 days to induce endotoxin tolerance [27] or orthotopic inoculation of 4T1 aggressive breast tumor cells. Scaffold implants and major organs were collected from LPS-tolerant mice one day after the final LPS dose or when 4T1 primary tumors reached ∼1,000 mm^3^. Tissues from healthy mice served as the control. The collected tissues were processed into single-cell suspensions and analyzed by flow cytometry to characterize immune-cell composition and phenotypes.

Repeated LPS exposure induced widespread immunosuppressive changes across major organs. Specifically, the overall frequency of CD4⁺ T helper cells (Th; CD45⁺CD3⁺CD4⁺) decreased, accompanied by increased frequencies of IL-4⁺ Th2 cells and CD25⁺Foxp3⁺ Tregs, and reduced frequencies of IFN-γ⁺ Th1 and IL-17A⁺ Th17 cells (**Figure. 1A**). LPS-tolerant mice also exhibited increased frequencies of Gr-1⁺ myeloid cells (CD45⁺CD11b⁺Gr-1⁺) compared with healthy controls (**Figure. 1B**). This population included Ly6G⁺ neutrophils and Ly6G⁻ monocyte-like cells, both of which expanded across major organs following repeated LPS exposure (**Figures. 1C and 1D**). Furthermore, LPS tolerance shifted macrophages and neutrophils from IL-12/IL-23 p40⁺ pro-inflammatory phenotypes toward IL-10⁺ anti-inflammatory phenotypes, while reducing the frequency of mature dendritic cells (DCs; CD86⁺MHC-II⁺) across major organs (**Figures. 1C-1F**). Together, these changes are consistent with the characteristic immune alterations associated with endotoxin tolerance [17; 18]. In contrast to the widespread immunosuppressive changes observed in major organs, implanted scaffolds maintained a distinct, immune-active environment. Following repeated LPS exposure, the frequencies of Th1 cells, Th17 cells, mature dendritic cells, IL-12/IL-23 p40⁺ neutrophils, IL-12/IL-23 p40⁺ monocyte-like cells, IL-12/IL-23 p40⁺ macrophages, and mature dendritic cells increased within the scaffolds. Conversely, the frequencies of Tregs and IL-10⁺ myeloid cells decreased. These changes contrasted with those observed in the corresponding pro-inflammatory and immunosuppressive cell populations in major organs.

**Figure 1.**
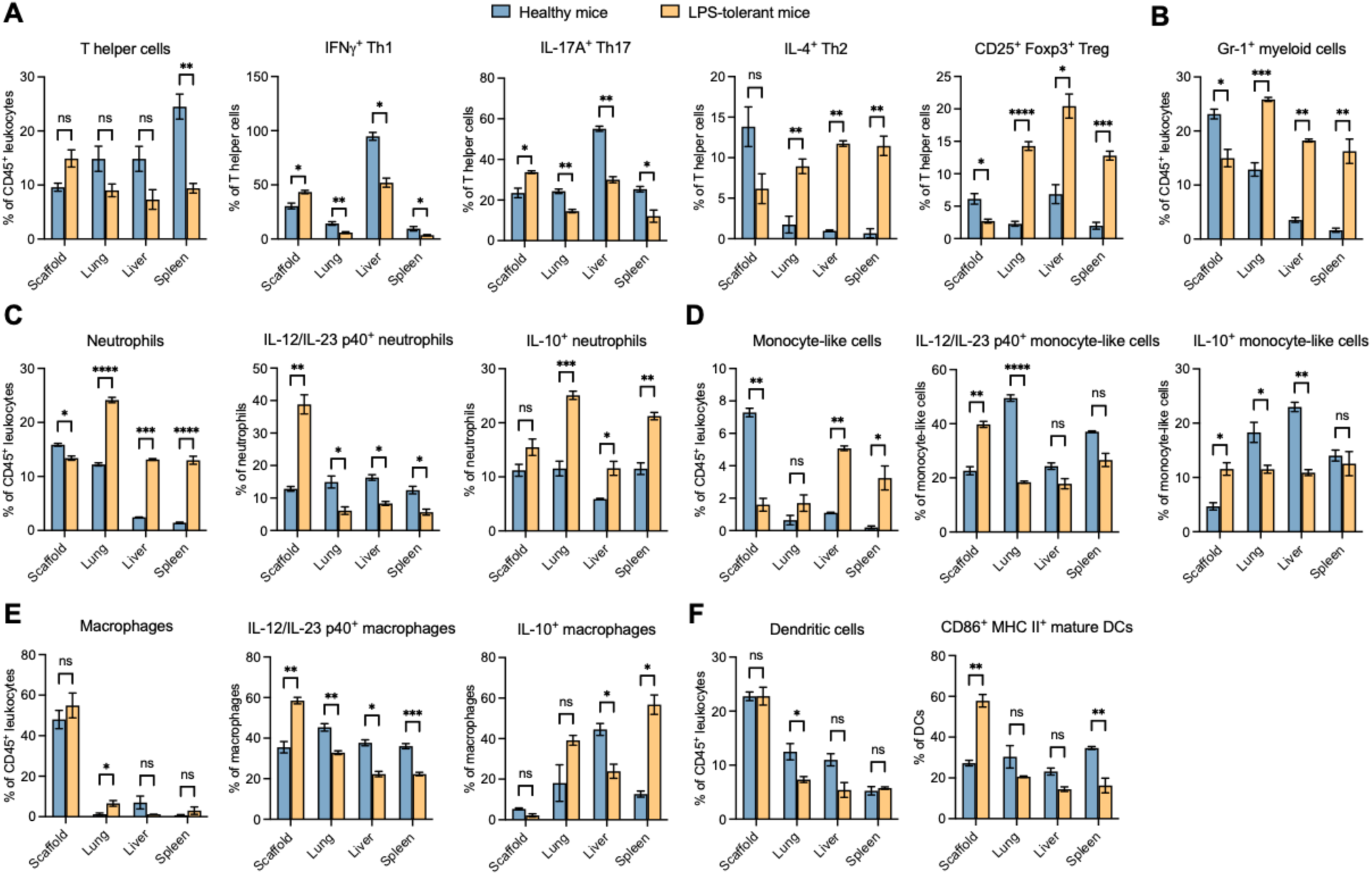
Immune cell composition and phenotypes in scaffolds, lung, liver, and spleen of healthy or LPS-tolerant BALB/c mice. **(A)** T helper cells and their subsets. **(B–D)** Gr-1⁺ myeloid cells, including neutrophils and monocyte-like cells, and their respective subsets. **(E)** Macrophages and their subsets. **(F)** Dendritic cells and their subsets. Frequencies of T helper cells (CD3⁺CD4⁺), Gr-1^+^ myeloid cells (CD11b⁺Gr-1⁺), neutrophils (CD11b⁺Gr-1⁺Ly6G⁺), monocyte-like cells (CD11b⁺Gr-1⁺Ly6G⁻), macrophages (CD11b⁺F4/80⁺), and DCs (CD11c⁺F4/80⁻) were quantified among CD45⁺ leukocytes. Subsets of each immune cell population were further defined using additional markers. DCs, dendritic cells; ns, not significant. *p < 0.05, **p < 0.01, ***p < 0.001, and ****p < 0.0001 by Student’s *t*-test.

Taken together with our previous finding in 4T1 tumor-bearing mice, we concluded that implanted scaffolds preserve a distinct, pro-inflammatory immune environment under two mechanistically different forms of systemic immunosuppression: aggressive cancer and endotoxin tolerance. This suggests that the FBR sustains local immune activity despite systemic immune dysregulation, revealing a potentially important homeostatic property of host–biomaterial interactions.

### LPS tolerance and aggressive tumors induce scaffold-localized upregulation of pro-inflammatory mediators

Scaffolds and selected major organs were collected from healthy, LPS-tolerant, and 4T1 tumor-bearing BALB/c mice and processed to generate tissue-conditioned media for cytokine profiling using a cytokine array. In LPS-tolerant mice, both the lungs and liver were examined because these organs are major sites of systemic innate immune responses to LPS. The liver plays a central role in clearing circulating endotoxin and regulating inflammatory responses, whereas the lungs contain extensive vascular and immune-cell networks that are highly responsive to LPS-induced immune reprogramming [29]. In 4T1 tumor-bearing mice, the lungs were selected because they represent a primary site of spontaneous metastasis in this breast cancer model and are therefore particularly relevant to investigating cancer-induced immunosuppression and the formation of a metastasis-supportive environment [30].

In response to repeated LPS exposure, multiple pro-inflammatory mediators were significantly reduced in major organs, consistent with the characteristic immune alterations associated with endotoxin tolerance [17; 18]. Notably, CXCL1, a pro-inflammatory chemokine that signals through CXCR2 to recruit and activate neutrophils at sites of inflammation [31], was significantly decreased in both the lungs and liver. Additionally, CCL5 and IL-6 were reduced in the lungs, whereas CCL9, G-CSF, and TNF-α were reduced in the liver (**Figures. 2A and 2B**). Conversely, several anti-inflammatory and immunoregulatory cytokines were elevated in the lungs of LPS-tolerant mice, including the type 2-associated cytokines IL-4 and IL-9 and the immunosuppressive cytokine IL-10. However, these cytokines did not increase significantly in the liver, suggesting organ-specific differences in immune reprogramming following repeated LPS exposure. Together, these cytokine alterations indicate attenuated pro-inflammatory signaling in major organs of LPS-tolerant mice, consistent with the immune-cell phenotypes described above and supporting the establishment of organ-specific immunosuppressive environments.

**Figure 2.**
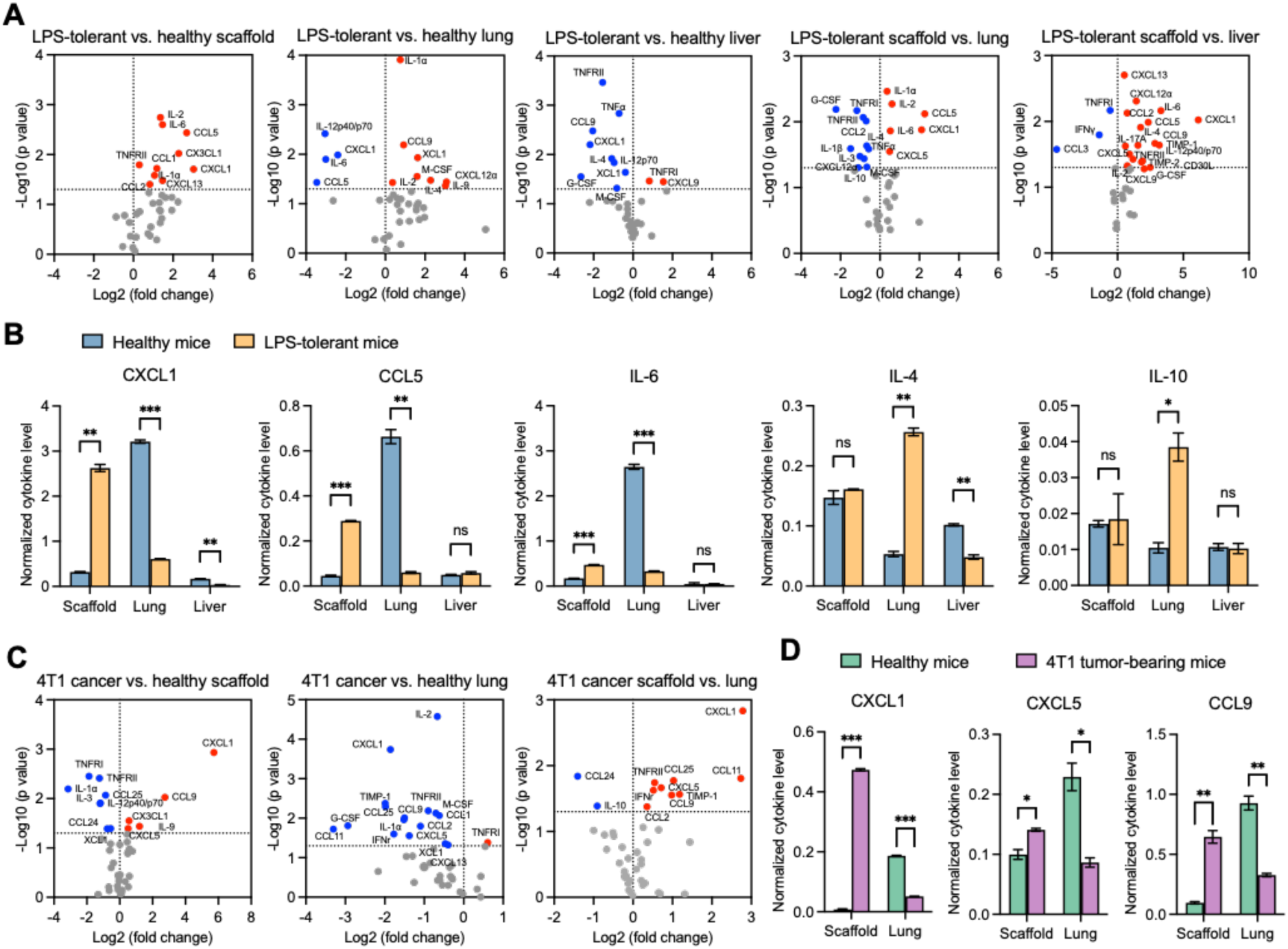
Cytokine profiles of tissue-conditioned media from implanted scaffolds and major organs of healthy, LPS-tolerant, and 4T1 tumor-bearing BALB/c mice. **(A,B)** Comparison of cytokine profiles and representative cytokine levels in conditioned media collected from implanted scaffolds, lungs, and livers of LPS-tolerant mice and their corresponding tissues from healthy controls. **(C, D)** Comparison of cytokine profiles and representative cytokine levels in conditioned media collected from implanted scaffolds and lungs of 4T1 tumor-bearing mice and their corresponding tissues from healthy controls. ns, not significant. *p < 0.05, **p < 0.01, ***p < 0.001, and ****p < 0.0001 by Student’s *t*-test.

In contrast, implanted scaffolds exhibited enhanced pro-inflammatory cytokine production following the induction of endotoxin tolerance. Compared with scaffolds from healthy mice, those from LPS-tolerant mice showed significantly elevated levels of CXCL1, CCL5, IL-6, and several other pro-inflammatory mediators, whereas IL-4 and IL-10 levels remained largely unchanged (**Figures. 2A and 2B**). Consequently, scaffolds maintained substantially higher levels of pro-inflammatory mediators than the corresponding major organs under systemic immunosuppressive conditions. A similar divergence was observed in 4T1 tumor-bearing mice, in which CXCL1 and other pro-inflammatory chemokines, including CXCL5 and CCL9, increased within implanted scaffolds but decreased in the lungs during cancer-induced immunosuppression (**Figures. 2C and 2D**).

Collectively, these findings reveal distinct cytokine responses within implanted scaffolds and major organs under two mechanistically different forms of systemic immunosuppression. Comparisons between diseased scaffolds and diseased organs demonstrate that scaffolds retain a more immune-active environment despite systemic immunosuppression. Importantly, comparisons between scaffolds from healthy and diseased mice further reveal that the scaffold-associated FBR is not merely preserved but undergoes local cytokine remodeling in response to systemic pathological conditions. This remodeling, characterized by increased production of pro-inflammatory mediators without corresponding increases in key immunosuppressive cytokines, may counteract the suppressive influence of infiltrating disease-associated immune cells and contribute to maintaining local immune activity despite systemic immunosuppression.

### Decorin reprograms suppressive immune cells toward pro-inflammatory phenotypes

We next investigated the mechanisms by which the scaffold-associated FBR sustains local immune activity despite systemic immunosuppression. Decorin, an extracellular matrix proteoglycan that can function as a DAMP and activate TLR2/4 signaling [21; 22; 23], exhibited striking differences in abundance between implanted scaffolds and major organs. Under healthy conditions, decorin abundance within scaffolds was ∼2-fold and ∼3-fold higher than that in the lungs and liver, respectively. Following repeated LPS exposure or tumor inoculation, decorin abundance increased within scaffolds but markedly decreased in major organs, further amplifying these differences. In LPS-tolerant mice, decorin protein levels in scaffolds were ∼23-fold and ∼40-fold higher than those in the lungs and liver, respectively. Similarly, in 4T1 tumor-bearing mice, decorin abundance within scaffolds was ∼3-fold higher than that in the lungs. These contrasting expression patterns suggested that decorin may contribute to the preservation of FBR-associated local immune activity under systemic immunosuppressive conditions. We therefore hypothesized that elevated decorin within implanted scaffolds promotes the reprogramming of infiltrating disease-associated immune cells toward pro-inflammatory phenotypes, thereby counteracting local immunosuppressive influences and sustaining an immune-active environment at the FBR site.

To test this hypothesis, we isolated lung and liver tissues from LPS-tolerant mice and lung tissues from 4T1 tumor-bearing mice. These tissues were dissociated into single-cell suspensions and treated with recombinant decorin proteins for 24 h, followed by flow cytometric analysis of immune cell phenotypes. As hypothesized, decorin treatment shifted multiple myeloid cell populations, including neutrophils, monocyte-like cells, and macrophages, from IL-10⁺ immunosuppressive phenotypes toward IL-12/IL-23 p40⁺ pro-inflammatory phenotypes. Decorin treatment also markedly increased the frequency of mature dendritic cells (**Figure. 3C**, **3D**). These findings are consistent with previous studies identifying soluble decorin as an endogenous DAMP that modulates innate immune responses primarily through its effects on myeloid cells [21; 22; 23]. In addition to its effects on myeloid cells, decorin treatment altered T helper cell polarization, as evidenced by increased frequencies of Th1 and Th17 cells as well as a decreased frequency of Th2 cells (**Figures. 3C**, **3D**). Given that decorin is primarily recognized for its immunomodulatory effects on myeloid cells, these changes in T cell phenotypes may arise indirectly through interactions between T cells and decorin-activated myeloid cells. In particular, enhanced production of pro-inflammatory cytokines by decorin-activated myeloid cells may promote Th1 and Th17 polarization while limiting Th2 responses. Together, these findings suggest that decorin may influence both innate and adaptive immune responses, potentially through myeloid–lymphoid crosstalk, thereby contributing to the maintenance of an immune-active environment within implanted scaffolds under systemic immunosuppressive conditions.

**Figure 3.**
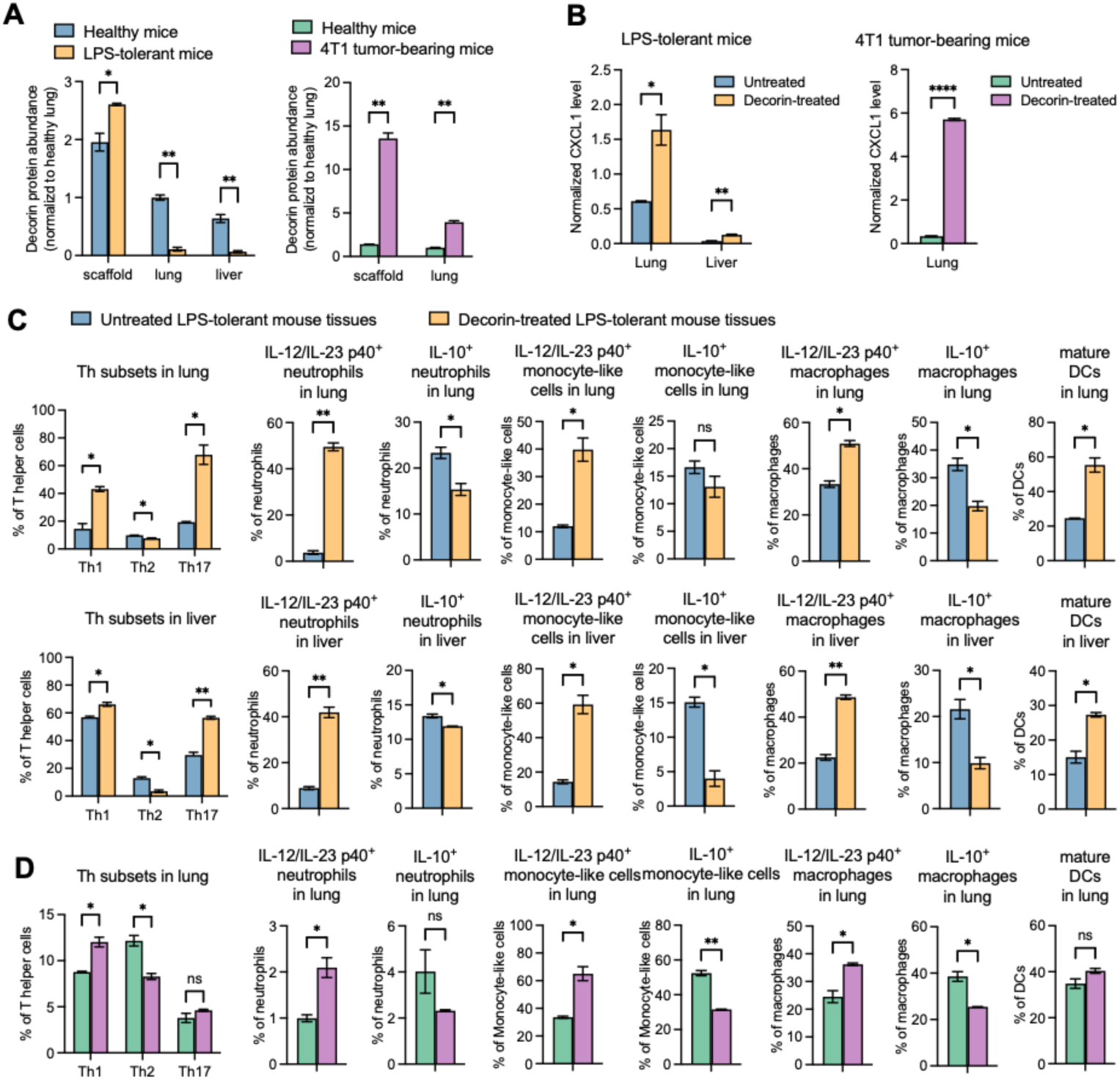
Decorin is upregulated in implanted scaffolds under systemic immunosuppression and promotes pro-inflammatory reprogramming and CXCL1 secretion by disease-associated immune cells. **(A)** Decorin protein abundance in scaffolds and major organs collected from healthy, LPS-tolerant, and 4T1 tumor-bearing BALB/c mice. **(B–D)** Recombinant decorin promotes CXCL1 secretion (B) and shifts disease-associated immune cells toward pro-inflammatory phenotypes (C, D) in cells isolated from LPS-tolerant and 4T1 tumor-bearing mice. *p < 0.05, **p < 0.01, ***p < 0.001, and ****p < 0.0001 by Student’s *t*-test.

While previous studies have primarily investigated the role of decorin in exacerbating inflammatory diseases, our findings reveal a potentially beneficial immunoregulatory function: reprogramming disease-associated suppressive myeloid cells toward pro-inflammatory phenotypes. This activity may contribute to the preservation of local immune function within the scaffold-associated FBR under systemic immunosuppressive conditions.

### Decorin promotes CXCL1 secretion to recruit pro-inflammatory immune cells

Cytokine profiling of scaffolds and major organs from healthy, LPS-tolerant, and 4T1 tumor-bearing mice revealed a consistent pattern of CXCL1 regulation. Following the induction of endotoxin tolerance or cancer-associated immunosuppression, CXCL1 abundance increased within implanted scaffolds but decreased in major organs, whereas changes in other cytokines varied across organs and disease models (**Figure. 2**). Given this consistent and contrasting pattern, we investigated whether decorin could stimulate CXCL1 production by cells derived from immunosuppressed organs. Single-cell suspensions prepared from the lungs and liver of LPS-tolerant mice and the lungs of 4T1 tumor-bearing mice were treated with recombinant decorin for 24 h. Compared with untreated controls, decorin treatment increased CXCL1 secretion 3–4-fold in lung- and liver-derived cells from LPS-tolerant mice and ∼17-fold in lung-derived cells from 4T1 tumor-bearing mice (**Figure. 3B**). These findings demonstrate that decorin enhances CXCL1 secretion by cells from immunosuppressed tissues, with the magnitude of the response varying between disease models. This variation may reflect differences in the composition or functional responsiveness of immune-cell populations associated with distinct immunosuppressive conditions.

CXCL1 is a potent chemokine that primarily recruits neutrophils through CXCR2-mediated chemotaxis, although additional immune cell populations may be recruited indirectly through secondary inflammatory signals generated by CXCL1-recruited cells [32]. We therefore hypothesized that elevated CXCL1 production within decorin-enriched scaffolds could promote the recruitment of pro-inflammatory immune cells, directly or indirectly, thereby contributing to the maintenance of an immune-active environment at the FBR site. To investigate this possibility, we performed a Transwell migration assay using splenocytes isolated from LPS-tolerant and 4T1 tumor-bearing mice. Single-cell suspensions of diseased splenocytes were prepared in serum-free medium and placed in the upper chambers of Boyden chambers, with recombinant CXCL1 added to the lower chambers as a chemoattractant. Following 24 h of incubation, cells that migrated into the lower chambers were collected and analyzed by flow cytometry. Their immune-cell composition and phenotypes were compared with those of the original, unsorted splenocyte populations.

Compared with unsorted splenocytes from LPS-tolerant mice, CXCL1-attracted cells exhibited a higher frequency of neutrophils (**Figure. 4A**). The migrated populations were also enriched in multiple pro-inflammatory immune-cell subsets, including IL-12/IL-23 p40⁺ neutrophils, IL-12/IL-23 p40⁺ monocyte-like cells, IL-12/IL-23 p40⁺ macrophages, and pro-inflammatory T helper cell subsets (**Figure. 4B**). Among these populations, neutrophils exhibited the most pronounced enrichment following CXCL1-directed migration. Similar patterns of neutrophil enrichment and increased representation of pro-inflammatory immune-cell subsets were observed among CXCL1-attracted splenocytes from 4T1 tumor-bearing mice (**Figures. 4C and 4D**).

**Figure 4.**
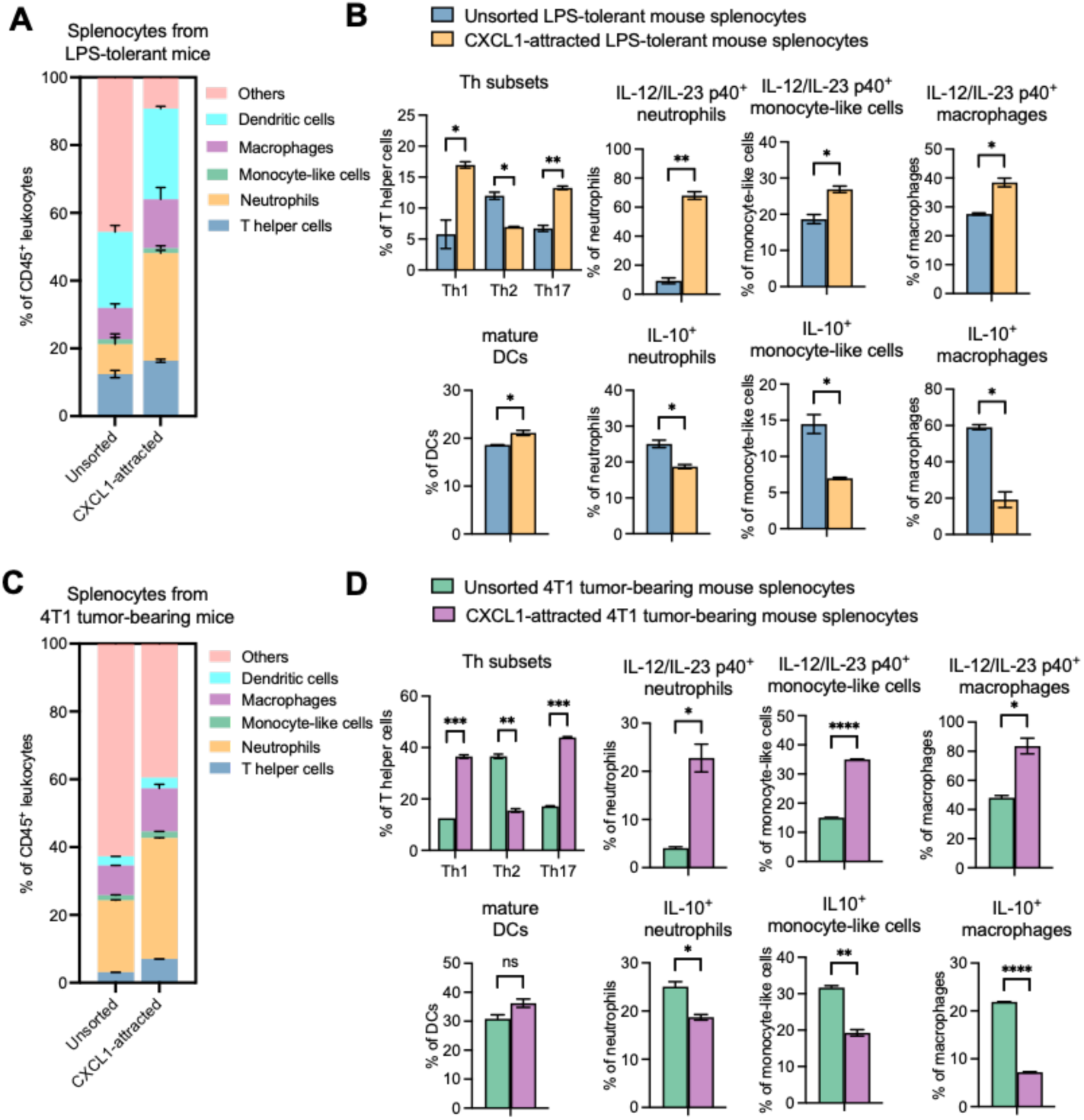
CXCL1 preferentially recruits neutrophils and enriches pro-inflammatory immune-cell populations from splenocytes of LPS-tolerant and 4T1 tumor-bearing mice. **(A)** Immune-cell composition and **(B)** phenotypic characterization of unsorted splenocytes and CXCL1-recruited splenocytes. ns, not significant. *p < 0.05, **p < 0.01, ***p < 0.001, and ****p < 0.0001 by Student’s *t*-test.

Collectively, these findings suggest that decorin-induced CXCL1 production may contribute to the maintenance of an immune-active environment within implanted scaffolds by promoting the recruitment and enrichment of pro-inflammatory immune cell populations. Together with the ability of decorin to shift disease-associated myeloid cells toward pro-inflammatory phenotypes, these results support a potential dual mechanism by which decorin-associated signaling sustains local immune activity at the FBR site: reprogramming infiltrating immune cells and promoting CXCL1-mediated recruitment of additional immune cells despite systemic immunosuppression.

### Decorin activates suppressive myeloid cells through TLR2/4-mediated signaling

Decorin has been reported to interact with TLR2 and TLR4 on myeloid cells, activating downstream MyD88-dependent signaling pathways, including NF-κB and MAPK pathways. These signaling pathways promote inflammatory activation and the production of pro-inflammatory mediators, including CXCL1, IL-12, and TNF-α [33]. To determine whether decorin exerts its immunostimulatory effects on disease-associated suppressive myeloid cells through TLR2/4 signaling, we isolated neutrophils from the immunosuppressive lungs of 4T1 tumor-bearing mice and treated them with recombinant decorin for 24 h in the presence or absence of TLR2/4 inhibition. Cells and conditioned media were subsequently analyzed by flow cytometry and cytokine array, respectively. Two approaches were used to inhibit TLR2/4 signaling: oxidized phospholipid OxPAPC, a pharmacological inhibitor of TLR2/4 signaling, and a combination of anti-TLR2 and anti-TLR4 blocking antibodies.

In the absence of TLR2/4 inhibition, decorin treatment markedly enhanced phosphorylation of p38 MAPK and NF-κB p65 in neutrophils, indicating activation of TLR2/4 downstream inflammatory signaling pathways (**Figure. 5A**). This response was accompanied by increased expression or secretion of several pro-inflammatory mediators, including IL-12/IL-23 p40, CXCL1, CXCL5, CCL5, TNF-α, M-CSF, and IL-1β (**Figure. 5A**, **5B**). Both OxPAPC and the combination of TLR2- and TLR4-blocking antibodies significantly attenuated decorin-induced TLR2/4 signaling activation and inflammatory mediator production. These findings suggest that decorin promotes the inflammatory activation of disease-associated neutrophils through TLR2/4-dependent signaling, providing mechanistic evidence for its immunomodulatory effects under systemic immunosuppressive conditions.

**Figure 5.**
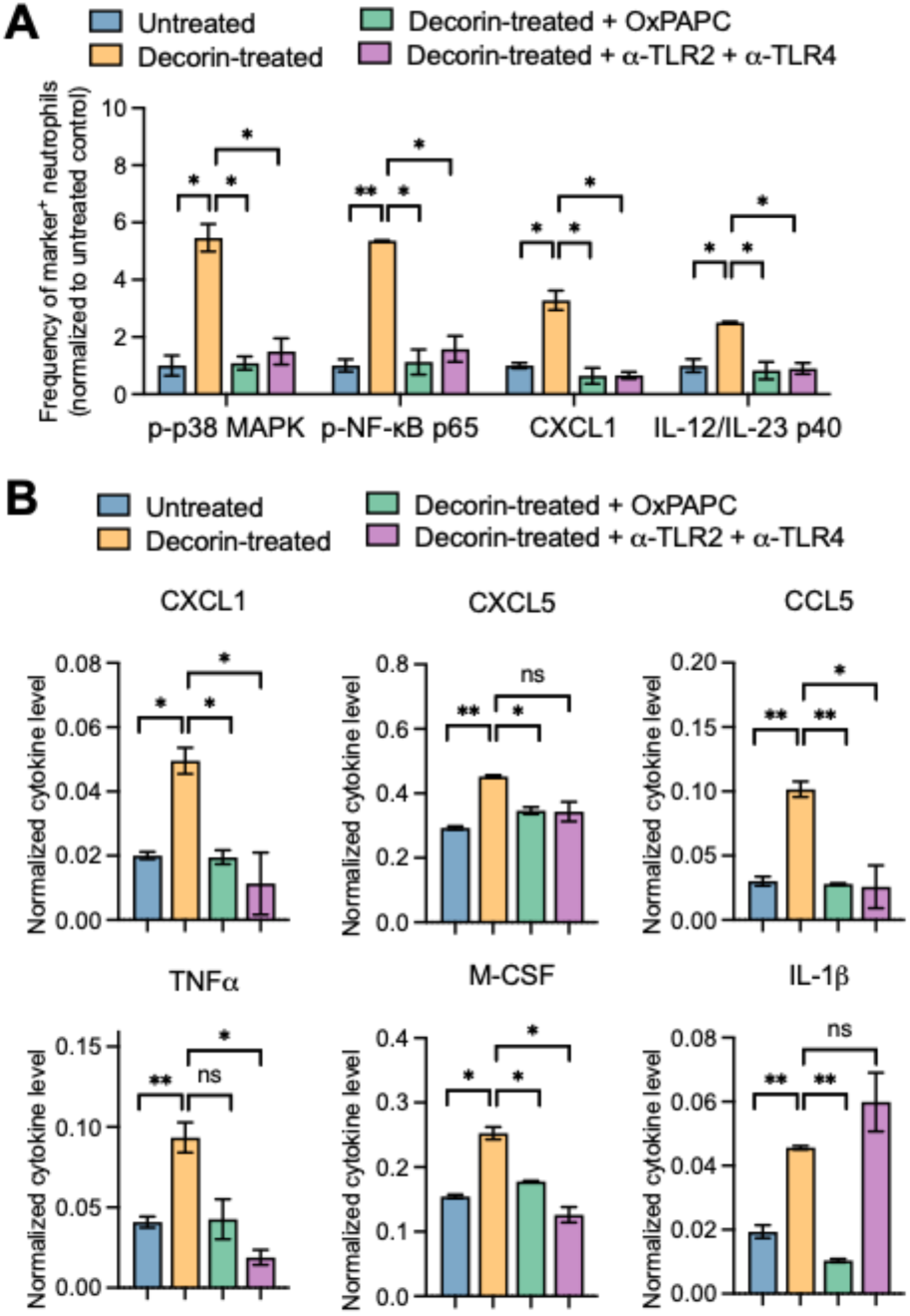
Decorin activates suppressive neutrophils and promotes pro-inflammatory cytokine production through TLR2/4 signaling. **(A)** Flow cytometric analysis and **(B)** cytokine profiling of Ly6G⁺ neutrophils treated with recombinant decorin in the presence or absence of TLR2/4 inhibitors. Neutrophils were isolated from the lungs of 4T1 tumor-bearing mice using magnetic-activated cell sorting (MACS). Isolated neutrophils were pretreated with OxPAPC or a combination of anti-TLR2 and anti-TLR4 blocking antibodies for 2 h, followed by incubation with recombinant decorin for 24 h. ns, not significant. *p < 0.05, **p < 0.01, ***p < 0.001, and ****p < 0.0001 by Student’s *t*-test.

Overall, this study reveals a previously underrecognized feature of the biomaterial-associated FBR: the ability to sustain localized immune activity despite systemic immunosuppression. We demonstrated this phenomenon in two mechanistically distinct models of systemic immune dysfunction: LPS-induced endotoxin tolerance and aggressive breast cancer. Mechanistically, our findings identify decorin, an extracellular matrix proteoglycan that also functions as a DAMP, as a potential mediator of this local immune response. Decorin abundance increased within implanted scaffolds under systemic immunosuppressive conditions, where it may counteract the suppressive influences of infiltrating disease-associated immune cells by promoting their reprogramming toward pro-inflammatory phenotypes. Furthermore, decorin enhanced CXCL1 production, which may facilitate the recruitment and enrichment of neutrophils and other immune-cell populations with pro-inflammatory phenotypes. These immunomodulatory effects were associated with TLR2/4-dependent activation of NF-κB and MAPK signaling pathways. Together, our findings suggest that decorin-associated immune signaling contributes to the preservation of an immune-active environment within biomaterial implants despite systemic immune dysfunction, revealing a potential mechanism through which the FBR adapts to pathological immune conditions.

## Discussion

FBR has traditionally been regarded as an undesirable host reaction to biomaterial implantation that can compromise implant functionality and therapeutic outcomes. Consequently, many biomaterial studies have sought to suppress the FBR because persistent inflammation, fibrosis, and encapsulation can impair drug release, biosensor performance, tissue integration, and the long-term function of implanted devices. [34; 35]. More recent studies, however, have increasingly explored how FBR-associated immune niches established within subcutaneous implants can be harnessed for disease monitoring, therapeutic immune modulation, and mechanistic investigations of disease progression. Porous synthetic scaffolds have been developed as biopsy-accessible immune niches that capture disseminated tumor cells, enabling early detection of metastatic disease and investigation of metastatic dissemination [36; 37; 38; 39; 40; 41]. Similar implantable immune niches have been applied in cancer and autoimmune disease models to collect innate immune cells whose phenotypes correlate with disease activity and therapeutic response, thereby establishing these implants as surrogate tissues for longitudinal monitoring of otherwise inaccessible organ sites [42; 43]. Biomaterial implants have also been engineered as active therapeutic niches. Infection-mimicking scaffolds containing GM-CSF, tumor antigens, and innate immune agonists recruit and program dendritic cells, producing antitumor immunity against poorly immunogenic tumors [44; 45; 46; 47; 48]. In regenerative medicine, biologically derived extracellular matrix scaffolds promote type 2 and immunoregulatory immune programs that support wound healing and constructive tissue repair. Within these niches, Th2 cells, macrophages, and specialized dendritic cell populations coordinate inflammation resolution, matrix remodeling, and tissue regeneration after injury, whereas synthetic materials elicit distinct immune states that may favor persistent inflammation or fibrosis [49; 50; 51; 52]. Scaffold implants can also provide controlled experimental tissues in which defined cellular, molecular, or matrix cues can be introduced to determine how local environments regulate disease progression. For example, co-implantation of melanoma cells with a drug-free urinary-bladder-matrix scaffold demonstrated that a scaffold-associated type 2 immune niche can suppress tumor establishment through CD4⁺ T cell- and macrophage-dependent mechanisms and enhance checkpoint blockade efficacy [53; 54]. Collectively, these studies redefine scaffold implants not simply as passive materials that must evade immunity, but as accessible engineered tissues for disease detection, longitudinal immune monitoring, therapeutic immune programming, tissue regeneration, and mechanistic investigation of disease progression.

Our findings introduce an additional consideration for these emerging applications: the FBR may actively maintain localized immune activity despite systemic immune dysfunction, potentially enhancing or interfering with the intended immunoregulatory functions of biomaterial implants. Many immunomodulatory biomaterials, including implantable scaffolds, are being developed for therapeutic applications in cancer, chronic inflammatory diseases, and other conditions characterized by immune dysregulation [55; 56; 57]. Therefore, understanding how systemic immune dysfunction influences local host responses to implanted biomaterials is essential for optimizing their immunomodulatory functions and therapeutic efficacy. In vaccine delivery, for example, the ability of the FBR to preserve localized immune activity under systemic immune dysfunction may offer an opportunity to improve vaccination in immunologically compromised populations, including patients with cancer and older adults experiencing immunosenescence. These populations often exhibit impaired innate immune responsiveness, dysfunctional antigen presentation, and weakened adaptive immune responses that can limit vaccine efficacy. Biomaterial-based vaccines provide a means of establishing localized immune niches that recruit and program antigen-presenting cells while controlling the delivery of antigens and adjuvants. Our findings suggest that the intrinsic immunoregulatory properties of the FBR could be harnessed to enhance the immunostimulatory functions of vaccine-delivery scaffolds. Previous studies have demonstrated that biomaterial-engineered immune environments can enhance vaccine efficacy by promoting immune-cell recruitment and activation [45; 46; 56; 58]. Our findings provide additional mechanistic insights into these effects, suggesting that decorin-associated TLR2/4 signaling and myeloid-cell reprogramming may contribute to the immunostimulatory properties of biomaterial-based vaccine platforms.

In tissue engineering, the ability of the FBR to sustain localized immune activity under systemic immune dysfunction may also have important implications. Anti-inflammatory drugs, such as dexamethasone, have been incorporated into biomaterial coatings and scaffolds to reduce inflammatory-cell recruitment, fibrotic encapsulation, and implant-associated tissue damage, thereby improving biocompatibility and device performance [59; 60; 61]. However, our findings suggest that FBR-associated immune responses may be influenced by the host’s pathological immune state, potentially affecting the efficacy of these anti-inflammatory strategies. In particular, enhanced decorin-associated TLR2/4 signaling under systemic immunosuppressive conditions may sustain local inflammatory activity and potentially counteract the intended effects of anti-inflammatory drug-releasing implants. Future studies should investigate whether local delivery of anti-inflammatory agents triggers compensatory inflammatory responses within the FBR environment in the diseased setting, given its apparent capacity to preserve local immune activity despite systemic immune suppression. If such compensatory responses occur, they may attenuate the therapeutic efficacy of anti-inflammatory drug-releasing implants. Overall, FBR-associated immune homeostasis may represent a previously underrecognized factor influencing the performance of anti-inflammatory implants, highlighting the importance of incorporating this consideration into rational biomaterial design and therapeutic evaluation, particularly in hosts with immune dysfunction.

Similarly, for biomaterial implants designed to deliver immunostimulatory or immunosuppressive therapeutics [62; 63; 64; 65], endogenous FBR-associated immune regulation may interact with the delivered agents and influence their therapeutic efficacy. In response to incorporated immunomodulatory agents, infiltrating disease-associated immune cells, and the biomaterials themselves, the FBR may dynamically regulate the balance between pro-inflammatory and anti-inflammatory signals, reshape local immune-cell phenotypes, and alter cytokine networks. These interactions may produce immune responses that differ from those predicted based on the pharmacological effects of the delivered agents alone. Thus, the FBR should be recognized as an active regulator of the local therapeutic environment rather than merely an adverse host response to implantation. Accordingly, the rational design of drug-eluting biomaterials should consider the intrinsic immunoregulatory properties of the FBR, the mechanisms of action of delivered therapeutics, and the pathological immune status of the recipient. Integrating these factors may enable more precise control of local immune responses and improve the predictability and therapeutic efficacy of implant-based interventions.

Beyond influencing implant performance, the disease-responsive immune activity of the FBR may provide a platform for identifying therapeutic molecules capable of counteracting immune dysfunction in major organs. Subcutaneously implanted scaffolds are accessible, experimentally controllable, and exposed to circulating immune cells and systemic mediators, yet their immune environments may respond differently from those of internal organs [12]. Comparing immune active scaffolds with immunosuppressed organs could therefore reveal endogenous molecules that are induced at the FBR site where immune balance is preserved but lost in diseased tissues. Candidate factors identified through such comparisons could subsequently be evaluated for their immunomodulatory effects in organ-derived cells or disease models, as demonstrated by our investigation of decorin in LPS-tolerant and 4T1 tumor-bearing mice. Decorin functions as an endogenous DAMP that activates TLR2/4-mediated, MyD88-dependent signaling in myeloid cells, promoting pro-inflammatory cytokine production [21; 22; 23]. Although previous studies have primarily investigated its roles in disease progression, fibrosis, tumor growth, extracellular matrix remodeling, and tissue repair [20; 21; 24], its potential as an immunostimulatory therapeutic remains relatively unexplored. Our findings suggest that decorin-mediated TLR2/4 activation may help restore inflammatory responsiveness in pathologically suppressed myeloid cells, highlighting decorin and potentially its bioactive fragments as candidates for developing immunostimulatory therapies targeting disease-associated immune dysfunction.

This framework may be particularly useful for developing personalized immunotherapeutic because immune dysfunction varies among diseases and among individuals with respect to the dominant suppressive cell populations, cytokine networks, disease stage, treatment history, and baseline immune competence [66; 67; 68]. Rather than assuming that one immune stimulant or anti-inflammatory agent will be appropriate for all settings, a scaffold-based immune niche could potentially help identify whether a particular host requires restoration of myeloid activation, inhibition of suppressive cytokines, enhancement of leukocyte recruitment, or resolution of excessive inflammation. This application remains conceptual and will require validation in clinically relevant models, but it positions the FBR as a potential functional platform for drug discovery and immune response stratification.

Future studies should determine whether FBR-associated immune homeostasis is conserved across additional forms of immune dysfunction, including treatment-induced immunosuppression, sepsis-associated immune paralysis, chronic infections, and autoimmune diseases. Longitudinal investigations are needed to establish when scaffold-associated immune environments respond to systemic immune dysregulation, which mechanisms sustain local immune activity across different pathological conditions, how long these responses persist, and how they influence FBR outcomes. It will also be important to determine whether biomaterials with different compositions, surface properties, and porous architectures employ shared or distinct mechanisms to preserve local immune homeostasis under systemic immune dysfunction. More importantly, future studies should investigate whether and how locally delivered immunostimulatory or immunosuppressive agents alter FBR-associated immune responses and potentially trigger compensatory mechanisms that maintain local immune activity. Understanding the interactions among the intrinsic immunoregulatory properties of the FBR, incorporated therapeutic agents, and the host’s pathological immune state will be essential for developing disease-adapted biomaterials. These insights may guide the rational design of immunomodulatory implants with more predictable immune responses and improved therapeutic efficacy across diverse biomedical applications.

Additionally, decorin belongs to the small leucine-rich proteoglycan (SLRP) family, whose members play diverse roles in extracellular matrix organization and cellular signaling. Beyond decorin, other SLRPs, including biglycan, lumican, fibromodulin, and osteoglycin, warrant investigation as potential regulators of scaffold-associated immunity [69; 70]. Although certain SLRPs can function as DAMPs and modulate innate immune signaling, their receptor interactions and immunoregulatory activities vary, and TLR2/4 agonism is not a universal property of this family. Decorin and other SLRPs have also been reported to antagonize TGF-β signaling and interact with inflammatory mediators, such as TNF-α, suggesting additional mechanisms through which they may regulate FBR-associated immune responses [22; 71; 72; 73]. Future studies should determine whether the FBR broadly harnesses multiple SLRPs to maintain local immune homeostasis or selectively engages specific family members depending on the disease context and host immune status. Elucidating how these SLRPs regulate infiltrating immune cells through distinct or overlapping mechanisms will be important for understanding the molecular basis of FBR-associated immune homeostasis.

## Author contributions

YZ conducted all the experiments and analysis. JW drafted the manuscript.

## Data availability

Data will be available on request.

## Ethics statement

All animal procedures were approved by the Institutional Animal Care and Use Committee (IACUC) at Iowa State University and were conducted in accordance with institutional and applicable federal guidelines.

## Acknowledgements

This work was funded by National Institutes of Health R21EB036758 (JW).

## Declaration of generative AI and AI-assisted technologies in the manuscript preparation process

During the preparation of this work, the authors used ChatGPT Plus for language editing. The authors reviewed and edited the output as needed and take full responsibility for the content of the published article.

